# Identification of key genes related to dexamethasone-resistance in acute lymphoblastic leukemia

**DOI:** 10.1101/337048

**Authors:** Qiuni Chen, Shixin Chen, Yuye Shi, Shandong Tao, Wei Chen, Chunling Wang, Liang Yu

## Abstract

Drug resistance is the main cause of poor chemotherapy response in acute leukemia. Despite the extensive use of dexamethasone(DEX) in the treatment of acute lymphoblastic leukemia for many years, the mechanisms of dexamethasone – resistance has not been fully understood. We choose GSE94302 from GEO database aiming to identify key genes that contribute to the DEX resistance in acute lymphoblastic leukemia. Differentially expressed gene(DEGs) are selected by using GEO2R tools. A total of 837 DEGs were picked out, including 472 up-regulated and 365 down-regulated DEGs. All the DEGs were underwent gene ontology(GO) analysis and Kyoto Encyclopedia of Gene and Genome(KEGG) pathway analysis. In addition, the DEGs-encoded protein-protein interaction (PPI) was screened by using Cytoscape and Search Tool for the Retrieval of Interacting Genes(STRING). Total 20 genes were found as key genes related to DEX resistance with high degree of connectivity, including *CDK1, PCNA, CCNB1, MYC, KPNA2, AURKA, NDC80, HSPA4, KIF11, UBE2C, PIK3CG, CD44, CD19, STAT1, DDX41, LYN, BCR, CD48, JAK1* and *ITGB1*. They could be used as biomarkers to identify the DEX-resistant acute lymphoblastic leukemia.

## Introduction

Acute lymphoblastic leukemia(ALL) is a heterogeneous hematologic malignant disorder characterized by the proliferation of immature lymphoid cells in the bone marrow, peripheral blood, and other organs. It is the most common childhood cancer and a major cause of death of adult leukemia^[1]^. The treatment of ALL comprises supportive therapy, anti-leukemia chemo-therapy, immunotherapy and other target therapy. The anti-leukemia chemo-therapy regime includes the administration of a glucocorticoid (prednisone, prednisolone, or dexamethasone), vincristine, and at least one other agent (usually asparaginase and anthracycline) ^[2]^. With the use of intensive chemotherapy regimen, the complete remission rates of adult ALL were 85 to 90%, but the long-term survival rates are only 30 to 50%^[3]^. Compared to the childhood ALL, the adult ALL patients have poorer prognosis, and the resistance to chemotherapeutic drugs, including dexamethasone(DEX) - resistance, is one of the most important cause. The mechanism of DEX–resistance was not fully understood and needed further investigation^[4]^.

Recently, gene expression profiling has been proven to be valuable in the pathogenesis, diagnosis and prognosis of many tumors including ALL^[5]^. Molecular analysis of the data from some public databases is a promising method to explore the disease related biomarkers. This article is aimed to identify the key genes and pathways relating to the DEX - resistant ALL patients, and it will be helpful to investigate the mechanisms of DEX resistance.

## Methods

### Microarray Data

GSE94302 from Gene Expression Omnibus(GEO) was chosen for this research, which is a public and freely available database (https://www.ncbi.nlm.nih.gov/geo/)^[6]^. GSE94302 is based on the following platform: GPL10558(Illumina HumanHT-12 V4.0 expression beadchip, Illumina Inc, San Diego, CA, USA). The preparation of samples was described in a public website (https://www.ncbi.nlm.nih.gov/geo/query/acc.cgi?acc=GSE94302). The dataset includes 99 samples, containing 58 DEX sensitive samples, 18 DEX resistant and 23 unknown ones. The known response samples were divided into two groups according to their regulation.

### Identification of differentially expressed genes (DEGs)

GEO2R(http://www.ncbi.nlm.nih.gov/geo/geo2r) was used to detect DEGs in sensitive and resistant samples. DEGs were defined as p< 0.05 and ∣logFC∣> 1 as the cut-off criterion. 837 DEGs were found, including 472 up-regulated and 365 down-regulated genes, and the top 10 genes with high degree of connectivity were chosen as key candidate genes in each group.

### Gene Ontology and Pathway Enrichment Analysis

The Gene Ontology(GO) concept was intended to use a common vocabulary to annotate the homologous gene and protein sequences in multiple organisms, thus we could query and retrieve genes and proteins based on their shared biology^[7]^. GO analysis and pathways analysis were carried out by KEGG PATHWAYS(http://www.kegg.jp/)^[8]^and DAVID(https://david.ncifcrf.gov/) ^[9]^ with p< 0.05 as the cut-off criterion.

### Protein-Protein Interaction (PPI) Network and Module Analysis

The online database STRING(https://string-db.org/) was used to develop DEGs-encoded proteins and protein-protein interaction network. Search Tool for the Retrieval of Interacting Genes (STRING) collects experimental and predictable information associated with the interactions of protein pairs in a given cell context via calculating the combined score of PPIs^[10]^. The DEGs were mapped into STRING, and the confidence score≥0.4, maximum number of interactors = 0 are set as the cut-off criterions. In addition, Cytoscape software^[11]^ was used to screen the modules of PPI network.

## Results

### Identification of DEGs and key genes

To identify the genes may be related to the dexamethasone sensitivity, DEGs between sensitive and resistant samples are screened using the GEO2R online analysis tool with p value < 0.05 and ∣logFC∣> 1. A total of 837 DEGs were identified, including 472 up-regulated and 365 down-regulated genes(Fig 1). Besides, 10 key genes in each group with high degree of connectivity were picked out (Table 1).

**Fig 1:**
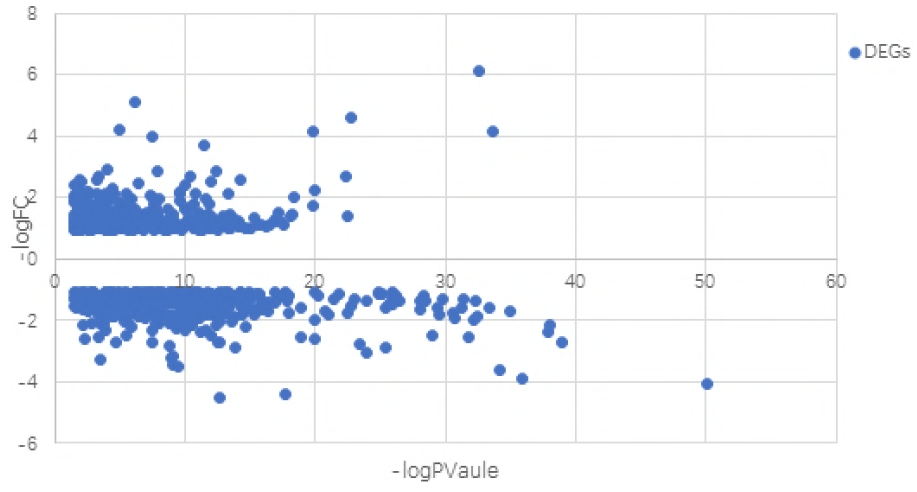
Distribution of DEGs. X-axis: −log p value, y-axis: −log∣FC(fold change)∣.DEGs were marked as blue dot.

**Table1:** TOP 10 key genes with higher degree of connectivity in two groups. A represents up-regulated DEGs. B represents down-regulated DEGs.

| Name | Degree | pvalue |
| --- | --- | --- |
| *CDK1* | 68 | 0.000000264 |
| *PCNA* | 60 | 0.0000076 |
| *CCNB1* | 56 | 2.07E-08 |
| *MYC* | 50 | 0.0000623 |
| *KPNA2* | 41 | 0.000777 |
| *AURKA* | 39 | 2.29E-08 |
| *NDC80* | 37 | 2.52E-14 |
| *HSPA4* | 36 | 7.54E-12 |
| *KIF11* | 34 | 3.11E-13 |
| *UBE2C* | 33 | 0.0000639 |

| Name | Degree | pvalue |
| --- | --- | --- |
| *PIK3CG* | 32 | 3.15E-08 |
| *CD44* | 29 | 0.00776 |
| *CD19* | 23 | 0.000117 |
| *STAT1* | 21 | 2.52E-11 |
| *DDX41* | 17 | 8.72E-17 |
| *LYN* | 17 | 0.0114 |
| *BCR* | 16 | 0.000804 |
| *CD48* | 14 | 1.24E-05 |
| *JAK1* | 13 | 1.35E-08 |
| *ITGB1* | 13 | 2.47E-12 |

### GO function and KEGG pathway enrichment analysis

In order to understanding the DEGs well, function and pathway enrichment analysis were conducted. DEGs were divided into two groups according to the criterion whether genes are up-regulated or down-regulated. All the up-regulated genes were put into DAVID software, the functions of up-regulated genes are screened (Fig 2A). The results showed that the up-regulated genes were particularly enriched in biological process(BP), including cell division, while in cell component(CC), they were enriched in following aspects: nucleus, cytosol, cytoplasm. For molecular function(MF), they were enriched in protein binding, poly(A) RNA binding (Table 2 A). The same method was used in the down-regulated genes. The functions of the down-regulated DGEs were shown in Fig 2B. These DGEs are enriched in BP, including interferon-gamma signaling pathway, cellular response to zinc ion and cadmium ion, negative regulation of growth, type I interferon signaling pathway. In terms of CC, the down-regulated genes were enriched in cell-cell adhering junction parts and membrane. For molecular function(MF), they were enriched in cadherin binding which involved in cell-cell adhesion (Table 2 B).

**Fig 2:**
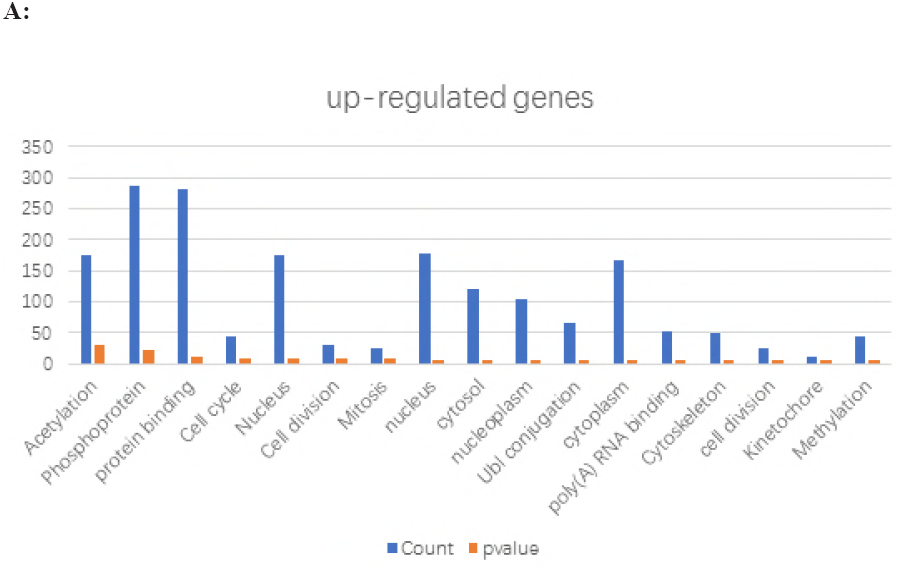
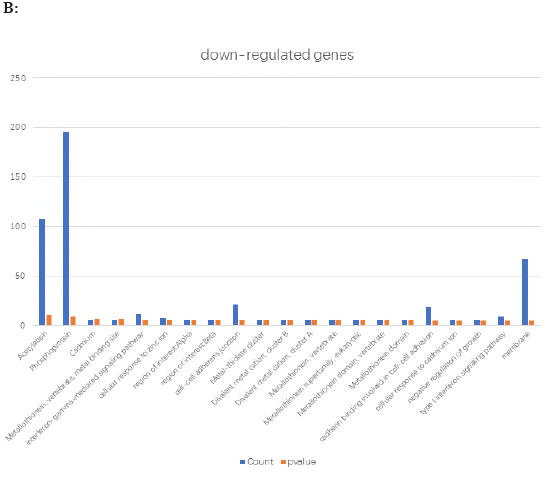
Gene ontology analysis of DEGs. A represents the functional annotation of up-regulated genes, and B represents the down-regulated genes.

**Table 2:** Gene ontology analysis of DEGs. A represents the up-regulated DEGs, and B represents the down-regulated DGEs. GO: Gene Ontology; FDR: False Discovery Rate; BP: biological process; CC: cell component; MF: molecular function

| Category | Term | Cou<br>nt | % | PValu<br>e | FDR |
| --- | --- | --- | --- | --- | --- |
| GOTERM_BP_DIR<br>ECT | **GO:0051301~cell**<br>**division** | 25 | 0.0343<br>26 | 6.49E-06 | 0.0112<br>77 |
| GOTERM_CC_DIR<br>ECT | **GO:0005634~nucleus** | 178 | 0.2443<br>98 | 1.19E-07 | 1.65E-04 |
| GOTERM_CC_DIR<br>ECT | **GO:0005829~cytosol** | 120 | 0.1647<br>63 | 4.41E-07 | 6.15E-04 |
| GOTERM_CC_DIR<br>ECT | **GO:0005654~nucleoplasm** | 103 | 0.1414<br>21 | 1.63E-06 | 0.0022<br>79 |
| GOTERM_CC_DIR<br>ECT | **GO:0005737~cytoplasm** | 166 | 0.2279<br>22 | 4.40E-06 | 0.0061<br>41 |
| GOTERM_MF_DIR<br>ECT | **GO:0005515~protein binding** | 283 | 0.3885<br>65 | 6.14E-13 | 9.04E-10 |
| GOTERM_MF_DIR<br>ECT | **GO:0044822~poly(A) RNA binding** | 53 | 0.0727<br>7 | 5.06E-06 | 0.0074<br>4 |

| Category | Term | Count | % | PValue | FDR |
| --- | --- | --- | --- | --- | --- |
| GOTERM_BP_DIR<br>RECT | **GO:0060333~interferon-gamma-mediated signaling pathway** | 11 | 0.0196<br>58 | 8.56E-07 | 0.0014<br>4 |
| GOTERM_BP_DIR<br>RECT | **GO:0071294~cellular response to zinc ion** | 7 | 0.0125<br>1 | 9.70E-07 | 0.0016<br>33 |
| GOTERM_BP_DIR<br>RECT | **GO:0071276~cellular response to cadmium ion** | 6 | 0.0107<br>23 | 1.21E-05 | 0.0203<br>93 |
| GOTERM_BP_DIR<br>RECT | **GO:0045926~negative regulation of growth** | 6 | 0.0107<br>23 | 2.21E-05 | 0.0371<br>42 |
| GOTERM_BP_DIR<br>RECT | **GO:0060337~type I interferon signaling pathway** | 9 | 0.0160<br>84 | 2.67E-05 | 0.0448<br>96 |
| GOTERM_CC_DIR<br>RECT | **GO:0005913~cell-cell adherens junction** | 21 | 0.0375<br>29 | 2.23E-06 | 0.0030<br>23 |
| GOTERM_CC_DIR<br>RECT | **GO:0016020~membrane** | 67 | 0.1197<br>35 | 3.02E-05 | 0.0409<br>95 |
| GOTERM_MF_DIR<br>RECT | **GO:0098641~cadherin binding involved in cell-cell adhesion** | 19 | 0.0339<br>55 | 6.40E-06 | 0.0092<br>67 |

represents the down-regulated DEGs.

In addition, KEGG pathway analysis was also employed. As it’s shown in Table 3, the up-regulated DEGs were enriched in cell cycle, biosynthesis of antibiotics, and p53 signaling pathway, while down-regulated DEGs were enriched in mineral absorption, platelet activation, hematopoietic cell lineage, actin cytoskeleton regulation, Epstein-Barr virus infection, graft-versus-host disease, peroxisome and fc gamma R-mediated phagocytosis signaling pathway.

**Table 3:**
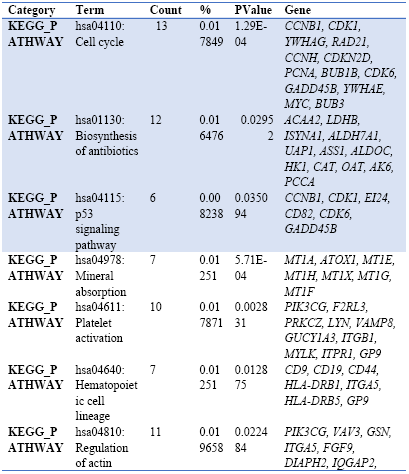
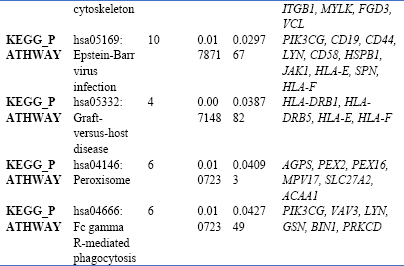
KEGG pathway analysis of DEGs. The up-regulated DEGs are marked in blue background, the others are down-regulated DEGs. KEGG: Kyoto Encyclopedia of Genes and Genomes

### PPI network and module analysis

Using the STRING online database and Cytoscape, the PPI network of the key genes from two groups(Table1) were made (shown in Fig 3). The key genes are selected by the criterion of higher connectivity. Based on the GO analysis and KEGG pathway analysis, the results show that the key genes from up-regulated gene group, including *CCNB1, CDK1, PCNA* and *MYC,* were associated with cell cycle pathway. For those from down-regulated DEGs, including *PIK3CG, CD19, CD44, LYN* and *JAK1,* were enriched in Epstein-Barr virus infection pathway.

**Fig 3:**
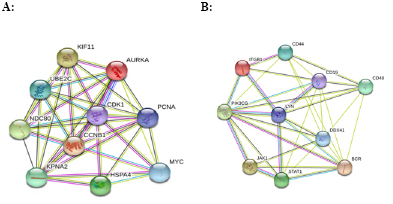
The protein-protein interaction network of DEGs. A: B: A: The protein-protein interaction network of top 10 hub gene in up-regulated DEGs. B represents the network of hub gens from down-regulated ones

## Discussion

Chemotherapy is the main treatment for pediatric and adult ALL, the therapeutic regimen contains induction, consolidation and maintenance therapy phrases ^[12]^. The induction therapy agents typically include glucocorticoid (prednisone or dexamethasone), vincristine and asparaginase, with or without anthracycline. According to previous studies, remission-induction treatment can eradicate the initial leukemic cell burden and restore the normal hematopoiesis (get complete remission, CR) in 96–99% of children and 78–92% of adult ALL^[13]^, but there were only 30% to 50% adult ALL patients achieve clinical cure^[3,14]^. The drug resistance is the main cause of the poor prognosis of adult ALL. Therefore, early prediction of the resistant-drug had an essential role in guiding ALL therapy.

Dexamethasone – resistance is the common cause of the poor prognosis of adult ALL. This study was focused on identifying the key genes related to dexamethasone - resistance in ALL, The GSE94302 from GEO database was chosen in the study. A total of 837 DEGs were picked up, including 472 up-regulated and 365 down-regulated genes. Then the DAVID software was used to analysis the DEGs in three aspects (MF, BP and CC). For deeper understanding, the up-and down-regulated genes are further clustered based on functions signaling pathways with significant enrichment analysis. The protein-protein interaction(PPI) network of DEGs is developed, and the top 10 genes were selected with the higher degree of connectivity.

The results shown that the up-regulated DEGs, including *CCNB1, CDK1, PCNA* and *MYC,* are biologically related to cell cycle regulation. Regulation of cell cycle has been proven to be closely related to tumor development and drug resistance^[15]^. Cohen Y et al. found that *CCNB1* was shown to be down-regulated in multiple myeloma and acute myeloid leukemia cells^[16]^. In this study, we found that high expression of *CCNB1* lead to ALL cells sensitive to dexamethasone treatment. *CDK1(* cyclin dependent kinase 1) is a member of the *Serine/Threonine* protein kinase family. Previous studies showed that *CDK1* was related to many types of tumors growth and the patients survival ^[17–21]^. Based on our analysis, *CDK1* was identified as the key gene in up-regulated DEGs, and GO annotations related to nucleoplasm, Ub1 conjunction cytoskeleton and cell division. The association of *CDK1* between drug resistance hasn’t been well explained. Therefore, our results could be helpful to explore its role in drug resistance deeply. *CCNB1* and *CDK1* have been found to regulated drug sensitiveness not only through cell cycle pathway, but p53 pathway according to our analysis. Previous study also found that *PCNA* was associated with cancers development ^[22, 23]^ and patients prognosis ^[24, 25]^. This study found that *PCNA* could serve as a biomarker indicating dexamethasone sensitivity through cell cycle pathway based on our analysis. *MYC* gene was first discovered in Burkitt lymphoma patients, it has been proven to play key role in the development of many tumors^[26–28]^. In our study, the level of *MYC* expression was related to the sensitiveness of dexamethasone treatment, and *MYC* regulated the drug sensitiveness through cell cycle pathway. This result need to be further validated in future studies.

The other key genes in up-regulated DEGs except for those enriched in cell cycle, including *KPNA2, AURKA, NDC80, HSPA4, KIF11, UBE2C.* Based on GO analysis, it was found that *KPNA2, AURKA, NDC80, HSPA4,* and *UBE2C* are enriched in protein binding. Drug’s performance can be enhanced or reduced by protein binding^[29]^. Recent research has found *KPNA2* is significantly associated with tumor differentiation, tumor depth, lymph node metastasis, venous invasion, recurrence and clinical response through L-type amino acid transporter 1^[30]^. In addition, *KPNA2* has been proven to be a potential marker of prognosis and therapeutic sensitivity in colorectal cancer patients^[31]^. But its mechanism needs further investigation. Our study also indicated that *KPNA2* could be a biomarker for ALL prognosis. *AURKA* plays an important role in the development of pediatric glioblastoma (pGBM), and its inhibitor is considered as the effective therapy of pGBM ^[32]^. *AURKA*, is a type of Aurora kinases. Recently, several Au0072ora kinase inhibitors were being investigated as novel anticancer therapy in breast cancer^[33]^. The importance of *AURKA* in ALL has not been studied deeply, our analysis will provide a new angle for further investigation. *NDC80* is overexpressed in a variety of human cancers^[34]^, its GO analysis suggests it is functional in protein binding, cell cycle, nucleus and cell division. The value of *NDC80* in leukemia has not been reported in previous studies. *HSPA4, KIF11* and *UBE2C* all have relationships with many carcinomas^[35–37]^, but their roles in leukemia need to be further studied.

The down-regulated genes are dominantly related to Epstein-Barr virus infection, platelet activation, hematopoietic cell lineage and fc gamma R-mediated phagocytosis, for example, *PIK3CG, LYN, CD19* and *CD44*. *PIK3CG* encodes a protein that belongs to the pi3/pi4-kinase family of proteins. The product of it is an enzyme which is thought to play a pivotal role in the regulation of cytotoxicity in NK cells. *PIK3CG* has been proven contributed to B cell development and maintenance, transformation, and proliferation^[38]^. The finding in this study indicted that *PIK3CG* regulated DEX sensitiveness through platelet activation, regulation of actin cytoskeleton, Epstein-Barr virus infection, and fc gamma R-mediated phagocytosis pathway. *LYN* has been a biomarker in many cancers^[39, 40]^. In this study, *LYN* also could be a sign on the response to dexamethasone. *CD19* is located on the surface of B lineage cells, except for plasma cells and on follicular dendritic cells. It has been used to diagnose cancers that arise from this type of cell - notably B-cell lymphomas^[41]^. *CD19* has also been proven to be a useful treatment target of B cell malignance. *CD19* may regulate drug resistance in ALL by hematopoietic cell lineage and Epstein-Barr virus infection pathway. *CD44* participates in a wide variety of cellular functions including lymphocyte activation, recirculation and homing, hematopoiesisand tumor metastasis. *CD44* expression is an indicative marker for effector-memory T-cells. The high levels of the adhesion molecule *CD44* on leukemic cells were essential to generate leukemia^[42]^. *CD44* was considered as cancer stem cell-like marker^[43]^. However, the value upon drug resistance of *CD44* in neoplasms remains controversial^[44]^. Therefore, it is necessary to identify the exact role of it in cancers.

*STAT1, DDX41, BCR, CD48, JAK1* and *ITGB1* are related to phosphoprotein according to the GO annotation. *STAT1* is a member of the STAT protein family. In mammals, the *JAK/STAT* pathway is the principal signaling mechanism for a wide array of cytokines and growth factors. *STAT1* and *JAK1* participate in the biological process via participating in type I interferons binding. *STAT1* is a key regulatory gene in autoimmune diseases and metastatic melanoma^[45, 46]^. Studies found that *JAK/STAT* pathway genes may play roles in lymphomagenesis, but they still need further investigation^[47–49]^. It has been shown that germline mutations in *DDX41* gene in several leukemia families, and *DDX41* mutation could develop neoplasia through acquisition of another somatic mutation. The recognition of *DDX41* mutated cases may have implications for surveillance, assessment of prognosis, and the design of targeted therapies^[50]^. Breakpoint cluster region(*BCR*) is at the chromosome 9 breakpoint. Although the *BCR-ABL* fusion protein has been extensively studied, the function of the normal *BCR* gene product is not clear. In our analysis, *BCR* may take part in DEX resistance, but its specific mechanism is unclear. *CD48* is a B-lymphocyte activation marker, expressed on all peripheral blood lymphocytes (PBL) including T cells, B cells, Null cell and thymocytes ^[51–53]^. *CD48* is being investigated among other markers in research on disease markers, and our study will be useful for research. *ITGB1*(integrin beta1) is associated with adverse prognosis of prostate cancer through regulating Caveolin-1 (*CAV1*), *CAV1* was over-expressed in prostate cancer and predicts adverse prognosis^[54]^. *ITGB1* was indicated to regulated drug resistance via platelet activation and regulation of actin cytoskeleton pathway according to our study, the relationships between *ITGB1* and leukemia need further research.

## Conclusion

In conclusion, this study identified several potential molecular targets that might contribute to the DEX resistance in ALL, including *CDK1, PCNA, CCNB1, MYC, KPNA2, AURKA, NDC80, HSPA4, KIF11, UBE2C, PIK3CG, CD44, CD19, STAT1, DDX41, LYN, BCR, CD48, JAK1* and *ITGB1*, which may function via cell cycle pathway, platelet activation pathway, or Epstein-Barr virus infection pathway. However, the finding here should be taken prudently.

## Acknowledgements

We are grateful to the researchers who provided their data for this analysis, and it is our pleasure to acknowledge their contributions.

